# Microbial population dynamics and evolutionary outcomes under extreme energy-limitation

**DOI:** 10.1101/2021.01.25.428163

**Authors:** William R. Shoemaker, Stuart E. Jones, Mario E. Muscarella, Megan G. Behringer, Brent K. Lehmkuhl, Jay T. Lennon

**Affiliations:** Department of Biology, Indiana University, Bloomington, IN, 47405, USA; Department of Ecology and Evolutionary Biology, University of California, Los Angeles, CA 90095, USA; Department of Biological Sciences, University of Notre Dame, Notre Dame, IN 46556, USA; Institute of Arctic Biology, University of Alaska Fairbanks, AK 99775, USA; Department of Biology and Wildlife, University of Alaska Fairbanks, AK 99775, USA; Department of Biological Sciences, Vanderbilt University, Nashville, TN 37232, USA

## Abstract

As the most abundant and diverse form of life on Earth, microorganisms commonly inhabit energy-limited environments where cellular maintenance and growth is highly constrained. To gain insight into how microorganisms persist under such conditions, we derived demographic parameters from a diverse collection of bacteria by censusing 100 populations in a closed system for 1,000 days. All but one taxon survived prolonged resource scarcity, yielding estimated times-to-extinction ranging over four orders of magnitude from 10^0^ – 10^5^ years. These findings corroborate reports of long-lived bacteria that have been recovered from ancient environmental samples, while providing insight into mechanisms of persistence. Critically, we found that as death rates declined over time, lifespan was extended through the scavenging of dead cells. Although growth and reproduction were dramatically suppressed in the absence of an exogenous resource supply, bacterial populations continued to evolve. Hundreds of mutations were acquired, contributing to genome-wide signatures of negative selection as well as molecular signals of adaptation. Remarkable consistency in the ecological and evolutionary dynamics indicate that distantly related bacteria respond to energy-limitation in a similar and predictable manner, which likely contributes to the stability and robustness of microbial life.

## INTRODUCTION

Microorganisms are the most abundant and diverse group of organisms on the planet. Their capacity for rapid growth across a range of conditions helps catalyze and regulate the biogeochemical processes that sustain life on Earth. Yet, microorganisms are often challenged by energy supplies that barely meet their basal metabolic demands^1–5^. As a result, many microorganisms in nature rest on the cusp of life and death^1,6–9^. Despite being recognized as a universal constraint, population and evolutionary dynamics that emerge under thermodynamically unfavorable conditions are rarely examined. Identifying the mechanisms through which this universal constraint affects microbial dynamics has the potential reshape our understanding of complex microbiomes in environmental, engineered, and host-associated ecosystems.

Across the tree of life, organisms are confronted with how to best allocate energy in order to maintain homeostasis when resources are scarce. Microorganisms have evolved a wide array of traits that can extend their lifespan and maximize fitness in energy-limited environments^3,10^. While some taxa have the flexibility to generate ATP using different combinations of electron acceptors and electron donors, other organisms are specialized in the uptake or assimilation of unique substrates^11–15^. Ultimately, variation in traits that contribute to persistence across taxa and the degree that they increase the lifespan of a typical microorganism is relatively unknown, even though energy limitation has important implications for the diversity, stability, and longevity of microbial systems^13,16–19^.

When energy replenishment is rare, energy-limited habitats can take on the properties of a closed system^20^. Under such conditions, dispersal is restricted and resource supply rates are governed by endogenous dynamics that involve the internal recycling of energy^21–23^. For example, dead cells can be scavenged by surviving individuals to meet their maintenance requirements and even support growth-related processes^24^. While there are likely thermodynamic limits, the extent to which necromass can sustain biological systems is unclear. Demographic properties such as population size and per capita death rate dictate necromass pool size and flux, which will inevitably diminish over time in closed systems, a consequence of necromass consumption and metabolic inefficiencies^25^. Fundamentally, these biophysical constraints place hard limits on the persistence and evolution of populations in a closed system. Nevertheless, genetic and phenotypic changes arise in model organisms exposed to prolonged starvation^26,27^, while selection at the molecular level has been documented in the “deep biosphere” where microbial biomass in the marine subsurface turns over on a time scale of hundreds to thousands of years, owing to extreme energy limitation^28^.

Here, using a combination of comparative and experimental approaches, we characterized the ecological and evolutionary dynamics of energy-limited bacteria. First, we used survival analysis to test demographic predictions while estimating times-to-extinction for a diverse collection of heterotrophic bacteria over 1,000 days. Second, we evaluated variation in demographic parameters among taxa and tested for predicted trade-offs using gene knock outs, physiological assays, and metabolic capacities inferred from whole-genome sequencing. Third, we tested whether longevity could arise from growth supported by cellular recycling using single-cell assays and metabolomic methods. Finally, to evaluate the extent to which bacteria evolve under energy limitation, estimating the frequency of *de novo* mutations and the strength of negative selection while testing for signatures of adaptation at the molecular level. Together, this multi-faceted approach allowed us to identify the predictable ways in which phylogenetically distant taxa survive and evolve in energy-limited environments.

## RESULTS AND DISCUSSION

All replicate populations survived ~1,000 days with zero exogenous resources in all but one taxon. Population dynamics were modeled using the Weibull distribution (Eq. 1), a distribution that allows for the net rate of growth to change over time and reduces to an exponential if the net rate of growth is constant. The survival curve deviated from first-order decay (*k* = 1) in over 90% of the populations. In all of these populations we found *k* < 1, indicating that the net growth rate (births - deaths) consistently increased over time (Fig. 1). Using the initial population size (*N*(0)), we found that the initial rate of cell deaths (*d*_0_ · *N*(0)) and the change in the net rate of growth (*k*) exhibited a strong negative relationship (Fig. 2), consistent with the interpretation that cells use necromass to meet maintenance energy requirements. Metabolomics, microscopy, and pooled population sequencing support the conclusion that the increase in the net rate of growth is driven by a decrease in the rate of death, which allowed us to calculate longevity as the mean time to death of a cell 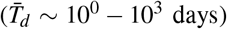 and the mean time to extinction of a population (*T_ext_* ~ 10^0^ – 10^5^ years) (Fig. 2).

**FIG. 1.**
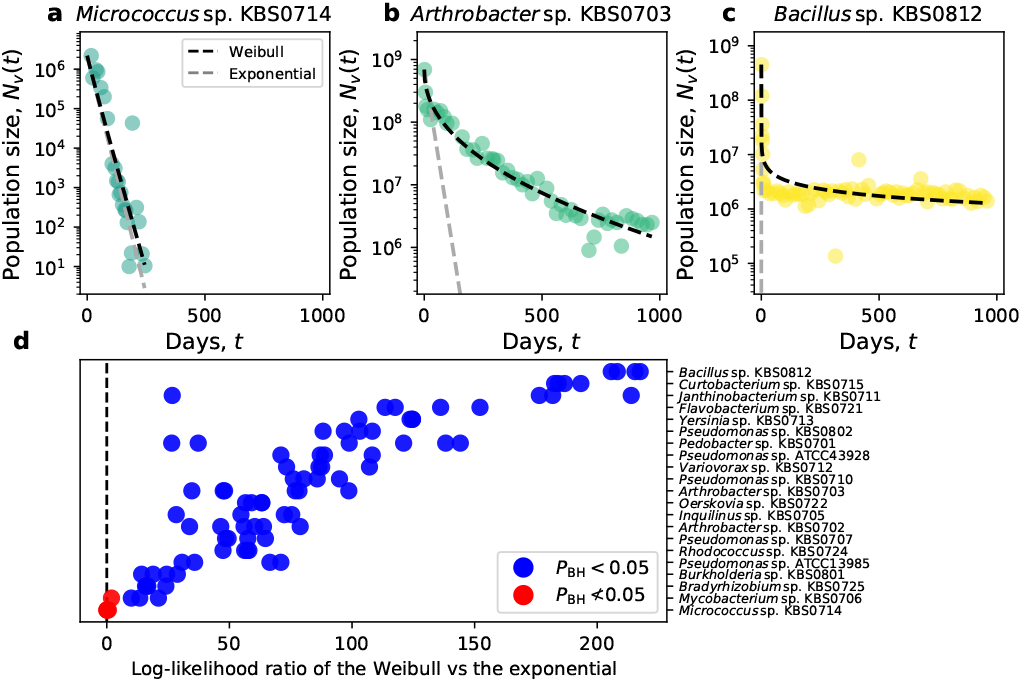
A high amount of variation was observed in the shape of the survival curves across species. The rate of decay of *Micrococcus* sp. KBS0714 (**a**) is almost linear on a semi-log scale, indicating that the net rate of growth does not change with time (*k* = 0.96 ≈ 1). While the survival curves of *Arthrobacter* sp. KBS0703 (**b**) and *Bacillus* sp. KBS0812 (**c**) exhibit clear curvature, meaning that the net growth rate is increasing over time. This curvature consistently occurs across taxa, as illustrated in **d** by the log-likelihood values of all replicate populations. We note the absence of phylogenetic signal in both parameters (Materials and Methods, SI Appendix, Fig. S9) (*LR* = 7.6, *λ_Pagel_ ≈* 0, *P* = 0.009).

**FIG. 2.**
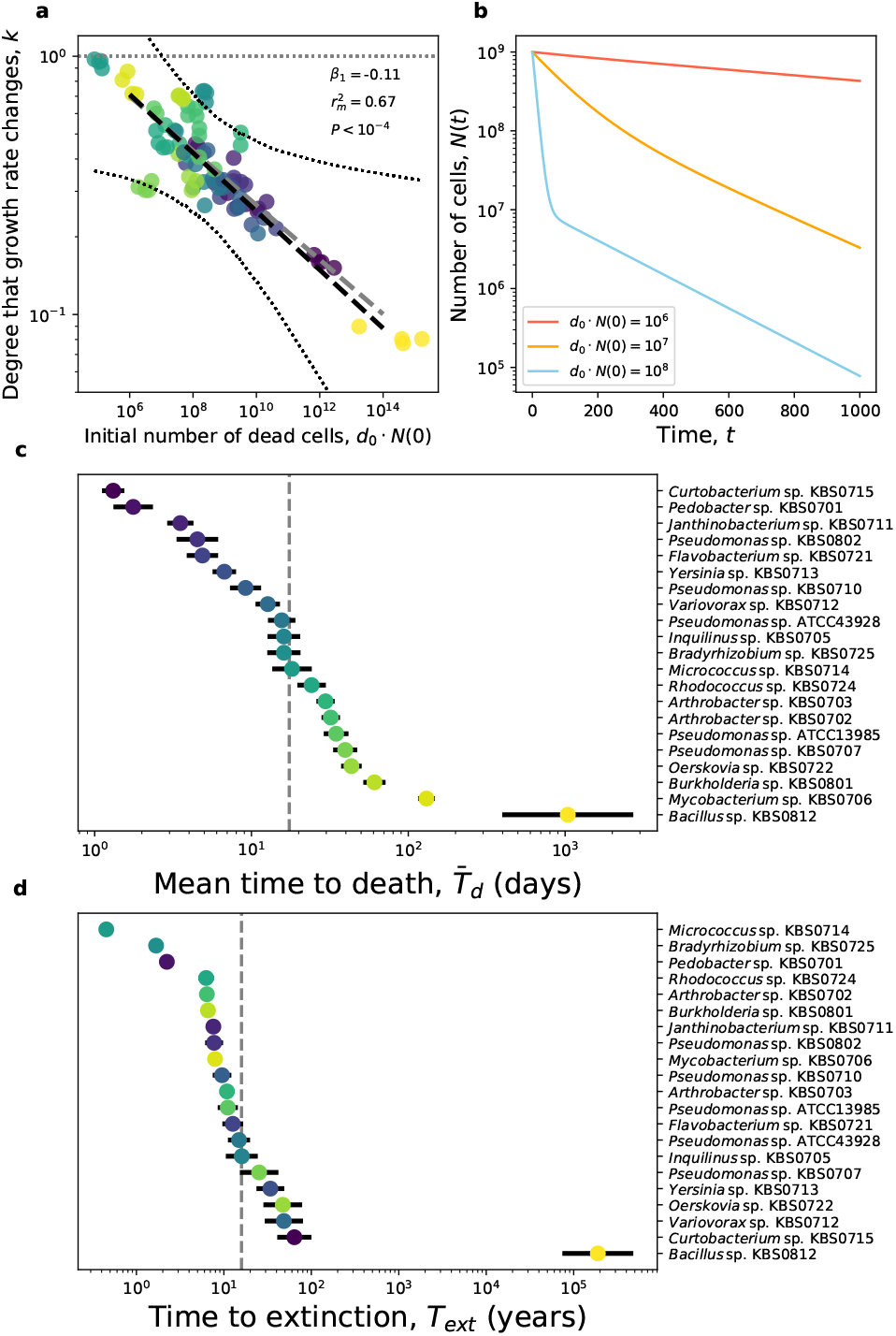
**a**: Populations with more deaths (larger *d*_0_ · *N*(0)) had a larger increase in the net rate of growth over time (smaller *k*), a prediction of necromass consumption models (Eq. 2) that is illustrated in **b**. **c, d:** Using these parameters, we examined the 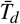 of an individual in each population and *T_ext_*. In **a**, dashed black and grey lines represent fixed effect and phylogenetic regressions, respectively. The dotted hull represents 95% confidence hulls. The dotted grey line represents the value of *k* where the net rate of growth does not change over time. Dots in **c** represent the mean 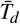 across replicate populations, bars represent twice the standard error, and the grey line represents the mean across all taxa.

### Growth rates increase over time under energy-limitation

The vast majority of microbial taxa exhibited survival curves where the net rate of growth increased over time (SI Appendix Fig. S1-S3). Though the amount of curvature in survival curves varied across taxa (Fig. 1), the only taxon with a survival curve resembling strict linear decay was *Micrococcus* sp. KBS0714. Unsurprisingly given the form of its survival curve, *Micrococcus* sp. KBS0714 was also the only taxon where all replicate populations went extinct, whereas all other taxa were still viable after 1,000 days. The curvature of survival curves on a semi-log scale has been previously reported^29–31^, but a comparative analysis of this phylogenetic breadth and timescale has not been performed.

We modelled this curvature using the Weibull distribution, an approach from survival analysis that is widely used in studies of demography, medicine, and engineering^32^. This distribution has two parameters, *d*_0_, the initial per-capita death rate and *k*, a shape parameter that describes how the net rate of growth changes over time (Eq. 1). Values of *k* greater or less than one reflect the degree that the net rate of growth increases or decrease over time, respectively, where the model reduces to exponential decay if *k* = 1. Model comparison of the Weibull and its reduced exponential form using a likelihoodratio test confirms visual inspection, as the Weibull distribution best explained the survival curves for all taxa except *Micrococcus* sp. KBS0714 (Fig. 1d). This deviation overwhelmingly occurred in one direction, as we find value of (*k* < 1 in all taxa that did not go extinct (SI Dataset S1). This bias indicates that the net rate of growth increased over time in energylimited environments across evolutionary distant taxa.

### Cellular recycling explains the increase in growth rate

Given that the net rate of growth increased over time for virtually all taxa, it is reasonable to suspect that the same underlying mechanism drove this pattern. In closed biological systems, the ability for microorganisms to meet maintenance energy requirements and generate new biomass likely depends on the one available resource: necromass. The initial flux of necromass (*S*) into the system should be proportional to the initial number of dead cells, defined as the initial number of cells times the initial per-capita death rate (∂*S_t_* ∞ *d*_0_ · *N*(0)). If the increase in the net rate of growth is due to cells scavenging dead cells for energy, the parameter *k* should decrease as the initial number of dead cells increases. We found that this was the case, as there was strong evidence of a negative relationship between the two variables. (*t*_19_ = −7.85, *r*^2^ = 0.67, *P* < 10^−5^) (Fig. 2a). By setting taxon identity as a random effect and examining the random slopes in a mixed model, we found that this pattern was also consistently found at the taxon level (Fig. 2a). This linear relationship also explains why *Micrococcus* sp. KBS0714, the taxon with smallest *d*_0_ · *N*(0), was also the only taxon that went extinct.

To evaluate whether the pattern in Fig. 2a was due to necromass consumption, we investigated whether dead cells accumulated in our populations. While the change in the proportion of dead cells over 1,000 days varied across taxa, it was nowhere near the expected value of ~ 1 if dead cells had accumulated without decomposing over 1,000 days (Materials and Methods and SI Appendix; Fig. S4). However, decomposition does not explain the absence of dead cells, as the concentration of cellular metabolites in the filtered supernatant at the 1,000-day mark was undetectable in all of the five taxa we examined (Fig. S5) and the concentration of amino acids was below the detection threshold in *Bacillus* (Fig. S6). Given that the number of cell deaths after 1,000 days can be greater than 10^9^ (~ 10^8^/mL) in these populations, the lack of detectable metabolites is consistent with the interpretation that individuals are consuming necromass. In addition, we found a strong relationship between *k* and the length of time until a taxon can enter an exponential phase of growth in energy-replete environments (i.e., lag time) (*t*_19_ = 3.38, *r*^2^ = 0.37, *P* = 0.0.0013) (Fig. S7), meaning that taxa that had a larger increase in the net rate of growth over time under energy-limitation also had a shorter lag time. This relationship is predicted by kinetic models of cellular growth, where lag time increases if cells do not have sufficient resources to maintain autocatalytic processes^33–35^. Finally, the survival curve trends we consistently observe can be reproduced by a simple mathematical model that describes the change in the net rate of growth as a consequence of necromass uptake via Monod kinetics (Eq. 2; Fig. 2b).

### Increase in the net rate of growth is driven by a decrease in death rate

While the early dynamics are primarily driven by massive cellular death, survival curves alone cannot tell us whether the change we observed over time was due to a decrease in death rate or an increase in birth rate. It is difficult to quantify birth rates in energy-limited populations that are out of equilibrium with a low net rate of growth. Nevertheless, the following logic indicates birth events were rare. First, there was no relationship between a sequence-based proxy of birth rate (the index of replication)^36^ and *k* (Fig. S8) (*t*_13_ = 0.69, *r*^2^ = 0.036, *P* = 0.500). Second, there was no relationship between the maximum growth rate (*μ_max_*) in energy-replete environments and *k*, which we would expect if the increase in the net rate of growth was primarily driven by a large increase in birth rate. Finally, the fact that no *de novo* mutations fixed over 1,000 days in any population indicates that very few generations occurred. This last observation allowed us to make some inferences about the number of birth events that occurred over 1,000 days.

By multiplying the maximum frequency of all *de novo* mutations in each replicate population by the final population size at day 1,000, we can obtain a proxy for the maximum size of a mutant lineage (*f_max_* · *N*(1,000) ≈ *N_mut_*; Table I). Because ≳ 95% of mutations are unique to a given population, we can assume that the majority of mutant lineages within a population arose after the initial culture was split into replicate populations. If lineages increased in size as a pure birth branching process, the number of generations required to generate a lineage of size *N_mut_* is log_2_(*N_mut_*). Using this assumption, we calculated a mean generation time of (log_2_(*N_mut_*)/1,000 d)^−1^ ≈ 60 – 75 days, three orders of magnitude larger than doubling time at the maximum rate of growth^37^. Continuing with our branching process assumption, we then calculated the minimum number of birth events (*b_min_*) necessary to generate *N_mut_* individuals and compared that estimate to *N*(1,000) with and without the curvature, providing an estimate for the contribution of birth events to the increase in *N*(*t*) due to necromass recycling. Because virtually all populations declined by several orders of magnitude in the first 100 days, we can say that the number of individuals at day 1,000 would be zero if the growth rate did not increase over time. This observation allowed us to estimate the minimum contribution of births to the difference in *N*(1,000) if the rate of growth did not increase over time as 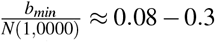 (Table 1). While this calculation is admittedly coarse, it suggests that birth cannot be the dominant demographic force.

**TABLE I.** Demographic estimates inferred from sequence data across taxa. The terms 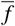 and *f_max_* represent the mean and maximum observed mutation frequency calculated from a mean of *n_mut_* mutations in a population, respectively. The term *b_min_* is the minimum number of cell divisions as a binary branching process calculated from *f_max_*. *T*_2_ is the mean time to cell division using *b_min_* and *b_min_/N*(1,000) is the percent that birth events contributed to the curvature at day 1,000.

| Taxon | $n_{mut}$ | $\bar{f}$ | $f_{max}$ | $b_{min}$ | $T_2$ days | $b_{min}/N(1,000)$ |
| --- | --- | --- | --- | --- | --- | --- |
| *Pseudomonas* sp. ATCC13985 | 208 | 0.14 | 0.64 | $7.3 \times 10^5$ | 50 | 0.30 |
| *Arthrobacter* sp. KBS0702 | 359 | 0.078 | 0.34 | $1.3 \times 10^5$ | 65 | 0.12 |
| *Curtobacterium* sp. KBS0715 | 79.3 | 0.14 | 0.56 | $1.2 \times 10^5$ | 66 | 0.20 |
| *Flavobacterium* sp. KBS0721 | 53.2 | 0.13 | 0.40 | $4.9 \times 10^4$ | 75 | 0.14 |
| *Oerskovia* sp. KBS0722 | 79.3 | 0.098 | 0.45 | $1.6 \times 10^6$ | 53 | 0.17 |
| *Rhodococcus* sp. KBS0724 | 53.0 | 0.11 | 0.32 | $8.9 \times 10^4$ | 69 | 0.088 |
| *Burkholderia* sp. KBS0801 | 29.0 | 0.14 | 0.41 | $5.5 \times 10^5$ | 55 | 0.15 |

### Longevity and extinction

Given that the dynamics we observed were primarily driven by a decrease in the rate of death, we used the Weibull parameters to calculate the mean time until a cell dies 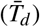 (i.e., lifespan). We found that 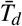 ranged from ~ 10^0^ – 10^3^ days (Fig. 2c), with a mean 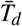 among all taxa of ~ 17 days and clear taxon-specific lifespans (*F*_20,73_ = 41.97, *P* < 10^−15^). These results suggest that a temporal decrease in the rate of growth is a common, but highly variable, response to energy limitation among phylogenetically diverse bacteria.

By specifying a lower observational threshold, we calculated the expected time until a given population was effectively extinct (Materials and Methods, SI Appendix). We found that *T_ext_* ~ 10^0^ – 10^3^ years for the majority of taxa, with *Bacillus* being an extreme case with a *T_ext_* of ~ 10^5^ years (Fig. 2d). These estimates of 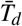 have the potential to inform ecological and evolutionary models that incorporate long-living pools of organisms with low metabolic activity (i.e., seed banks)^13,17,19^. While the timescales of the predicted values of *T_ext_* extend far beyond what can be experimentally observed, they may provide useful estimates for microbial dynamics that are difficult to observe in nature. Finally, the lack of extinction and high degree of comparability in population dynamics across phylogenetically distant taxa has the potential to inform the study of matter closed ecosystems (e.g., biospherics)^20^.

While Bacillus represented an extreme case due to its ability to form long-lasting protective endospores, its values of *N* · *d*_0_ and *k* do not deviate from the relationship we observed among all other taxa (Fig. 2a). This consistency was likely due to a large portion of Bacillus populations remaining in a vegetative state instead of forming spores (~ 70% vegetative cells), a finding that has been reported in experiments with different *Bacilli* strains^38,39^. However, this observation raises the question of, to what degree, does endospore formation increase 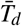 and *T_ext_* ? To answer this question, we constructed a *Bacillus* mutant that is incapable of forming spores and performed 80 day survival curves for the mutant (ΔspoIIE) and the WT (Materials and methods). WT and ΔspoIIE populations exhibited qualitatively similar survival curves, though the curve was consistently lower for ΔspoIIE. Despite the similarity in shape (Fig. S10), the removal of endospore formation had a dire effect on longevity, reducing 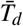 by 95% and *T_ext_* by 98%. While such reductions are substantial, the longevity of ΔspoIIE remained fairly high among the taxa we examined, suggesting that this singular life-history strategy is not the sole contributing factor for the longevity of Bacillus. Instead, it is likely that necromass recycling makes a nontrivial contribution to the longevity of *Bacillus* in energy limited environments.

### Evolutionary dynamics over 1,000 days

While the number of birth events is low, we attempted to investigate evolutionary dynamics from pooled population sequencing. We found that the distribution of mutation frequencies (i.e., the site frequency spectra) share a qualitatively similar form across taxa (Fig. S11), consistent with the similar population dynamics having occurred under energy limitation. Indeed, summary statistics indicate that the spectra are highly skewed across taxa, such that the difference between the mean number of pairwise differences and the number of segregating sites (Tajima’s D, *D_T_*) is consistently greater than zero for all taxa with a sufficient number of mutations (SI Appendix; Fig. S12). This pattern is consistent with a strong bottlenecking event due to a rapid reduction in population size. Indeed, *k* is correlated with *D_T_* at a borderline significant level (*β*_1_ = −3.10, *P* = 0.052), suggesting that the observed values of *D_T_* are at least in-part driven by the demographic history of the system (Fig. S13). However, since all replicates of a given taxon started from a single clone and very little standing genetic variation was shared across taxa, there was initially very little diversity that could undergo a bottleneck during the initial reduction in population size. This seemingly contradictive result can be resolved by considering an alternative demographic model, one where mutations initially increased in frequency, only to contract once necromass was depleted.

Despite the strong bottlenecking that likely occurred due to the initial rapid decline in population size, there was evidence that natural selection operated over the course of 1,000 days. The ratio of genome-wide nonsynonymous to synonymous polymorphic mutations (*pN/pS*) is less than one in the majority of taxa (Fig. 3), a signal of negative selection that has been found in microbial populations in the deep biosphere^26,27^. We then searched for potential targets of positive selection by identifying the set of genes that acquired more nonsynonymous mutations than expected by chance. We found that non-synonymous mutations were non-randomly distributed across genes in all taxa examined, a potential signal of adaptive evolution (SI Dataset S2, Fig. S14). By mapping genes to their higher-level functions, we identified the cellular features that contributed to adaptation across multiple taxa (i.e., convergent evolution). Three pathways were enriched for nonsynonymous mutations in two out seven taxa: lysine biosynthesis (M00016), pyrimidine biosynthesis (M00053), and branched-chain amino acid transport (M00237). However, while the lack of fixation events was a benefit for demographic inference, it ultimately limits our ability to make concrete claims about adaptation under energy-limitation.

**FIG. 3.**
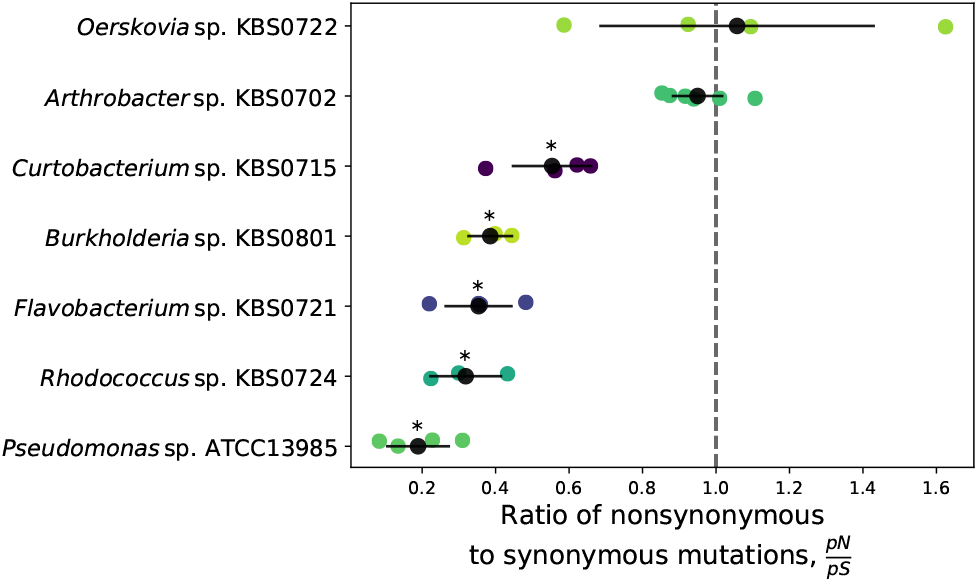
The ratio of nonsynonymous to synonymous mutations (*pN/pS*) within each taxon suggests that negative selection is occurring under extreme energy-limitation. The expected value in the absence of positive or negative selection is represented by a dashed grey vertical line. The black dot represents the mean within a given taxon across replicate populations and the black bars represent twice the standard error. Asterisks indicate that the mean is significantly less than one based off of a left-tailed one-sided *t*-test after false discovery rate correction.

## CONCLUSION

The work presented here constitutes one of the most rigorous estimates of bacterial longevity to-date, spanning over an extended length of time as well as across the tree of life. We estimated that bacterial cells can survive for ~ 10^0^ – 10^3^ days in the absence of exogenous resources, contributing to a population-level extinction timescale of ~ 10^0^ – 10^5^ years. We found that this longevity was primarily driven by the death rate decreasing over time due to necromass recycling. Though death was the predominant demographic force, a meager number of birth events occurred that generated hundreds of mutations, providing insight into the strength of selection and direction of molecular evolution under extreme energy-limitation. That necromass recycling was so prevalent among phylogenetically diverse taxa suggests that it may be a universal bacterial trait, allowing cells to survive extreme energy-limitation across dissimilar ecosystems.

## MATERIALS AND METHODS

### Constructing survival curves

We used a collection of bacterial strains with well-characterized functional traits, the majority of which were isolated from soil ecosystems and are described in detail elsewhere^37^. Single colony isolates of each taxon were inoculated into 50 mL Erlenmeyer flasks containing 20 mL of R2A medium. The flasks were incubated at 25 °C and shaken at 150 RPM. Before reaching stationary phase, cultures were split into 50 mL Falcon tubes at equal volumes and cells were pelleted in a centrifuged at 4,500 RPM. In order to remove residual medium, we discarded the supernatant and washed the cell pellet five times with pH 7.0 phosphate buffered saline (PBS). We initiated the survival curve experiments by dispensing the washed cells from each species into replicate (n = 4-8) 50 mL Falcon tubes, which were kept in the dark at 25 °C. We tracked the population density of each tube over time by plating dilutions onto R2A plates and counting the number of colony-forming units (CFUs). To allow for the emergence of slow growing individuals, we monitored colony formation on plates for up to two weeks (a typical strain formed colonies in < 48hrs.). The experiment was run until day 1,000, or until the population went extinct.

### Performing Survival Analysis

To statistically model the outcome where death rate decreases as a result of cells using resources supplied by dead cells, we used the survival function of the Weibull distribution:

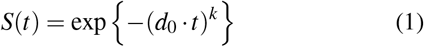

Where *d*_0_ is a scale parameter that represents the initial percapita death rate, *k* is a shape parameter that describes how the death rate of the system changes over time, and S(t) is the proportion of surviving individuals at time t. Because all populations are in environments where there zero initial exogenous resources, we can consider all initial changes in *N*(*t*) to be the result of death events. If *k* < 1 the death rate decreases over time, the opposite being true if *k* > 1. If *k* = 1 the death rate is constant with respect to time and population size decays exponentially. Model comparison was performed using the form of the likelihood ratio test provided by Wilks’ theorem. Using the estimated Weibull parameters, we calculated the mean time to death (i.e., lifespan) of a cell as 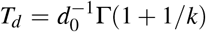, where Γ is the gamma function. The statistical relationship between *k* and *d*_0_ · *N*(0) was examined using a mixed-effects linear model via the lmer function (SI Appendix)^40^. The marginal coefficient of determination 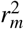 was estimated using the r.squaredGLMM function from the MuMIn package^41^. The confidence hull was estimated from a semi-parametric bootstrap using the bootMer function from lmer and phylogenetic linear regression using phylolm^42^ All survival analyses were conducted in R v3.5.0^43^.

### Necromass recycling model

We examine a system of equations that describe the dynamics of a system where viable cells (N) die at an initial death rate (*d*_0_) and enter a pool of dead cells (*N_d_*). These dead cells decay into a necromass carbon pool (S) at a given rate (δ) with a given amount of carbon per-cell (*c*). The uptake of necromass to meet per-cell maintenance energy requirements and increase the net rate of growth in terms of carbon (*m*) is described by Monod kinetics, where *V_max_* represents the maximum uptake rate of necromass and *K_S_* is the half-velocity constant.

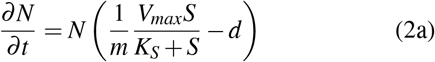

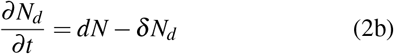

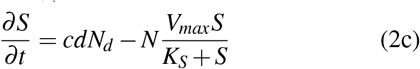

This system of equations models the net change in growth rate when maintenance energy costs can be met via the consumption of necromass.

### Mutation Calling

We performed pooled population sequencing on each tube with sufficient biomass. Biomass was concentrated at day 1,000 using MilliporeSigma™ Amicon™ Ultra Centrifugal Filter Units. Biomass pellets were then resuspended in 1mL PBS, centrifuged, and decanted before DNA was extracting using a MoBio UltraClean Microbial DNA Isolation Kit. Libraries were prepared using the NEXTflex™ Rapid DNA-Seq Kit (Bioo Scientific) and paired-end sequencing was performed using an Illumina NextSeq 500 High Output for 150 Cycles. Raw reads were cleaned and trimmed using cutadapt and mutations were mapped to hybrid *de novo* assembled Nanopore and Illumina genomes (see SI appendix for additional detail) and called with breseq v0.32.0^44,45^. We examined mutations in taxa with at least three replicates with a minimum mean genome-wide coverage of 50. Genome-wide likelihood ratios, *pN/pS*, and gene-specific multiplicity scores were calculated using publicly licensed code^46^. We tested whether *pN/pS* was less than one in each taxon using a left-tailed one-sided t-test and corrected for multiple testing using the Benjamini–Hochberg procedure from statsmodels^47^. The same test was used to determine whether *D_T_* was greater than one. Additional information on genetic diversity calculations can be found in SI appendix.

## AVAILABLE CODE AND DATA

We used open source computing code and custom scripts to complete our analyses. Raw reads used for assembly are available on SRA and assemblies are available on NCBI, with accession numbers listed in SI Dataset S4. The raw reads from evolved lines are available on SRA (BioProject PRJNA561216). All remaining data is available on Zenodo (doi:10.5281/zenodo.4458917). All analyses can be reproduced or modified for further exploration using code on GitHub: https://github.com/LennonLab/LTDE.

## ACKNOWLEDGMENTS

We thank the IUB CGB and the UNH HCGS for their help with the sequencing efforts. This work was supported by National Science Foundation Dimensions of Biodiversity Grant 1442246 and US Army Research Office Grant W911NF-14-1-0411. Genome assembly and mutation calling was supported by Lilly Endowment, Inc., through its support for the Indiana University Pervasive Technology Institute, the National Science Foundation under Grant No. CNS-0521433, and Shared University Research grants from IBM, Inc., to Indiana University.

