## Supplemental Information for "Microbial population dynamics and evolutionary outcomes under extreme energy-limitation"

#### This PDF file includes:

Supplementary text
Figs. S1 to S14
Legends for Dataset S1 to S4
SI References

#### Other supplementary materials for this manuscript include the following:

Datasets S1 to S4

### Supporting Information Text

**Survival Analysis.** We used the Weibull distribution to model survival curves. We assume that death events occur more frequently than birth events at any given time (i.e., $d(t) >> b(t)$) such that we can describe the system using equations do not increase at any point in time time (i.e., a monotonically decreasing function). The Weibull distribution is a two-parameter continuous distribution that is often used to model systems where the failure rate changes over time (1).

We start with the form of the Weibull that describes the number of cells at time t ($N(t)$):

$$N(t) = N(0) * \exp\left\{-(t \cdot d_0)^k\right\} \quad [1]$$

Where $d_0$ is a scale parameter that describes the spread of the distribution, k is a shape parameter that describes how the failure rate of the system changes over time, and $N(0)$ is the initial number of cells. If $k < 1$ the failure rate of the system decreases over time (i.e., death rate increases), the opposite being the case if $k > 1$. If $k = 1$ the failure rate remains constant through time and population size decays exponentially. After dividing both sides by $N(0)$, we are left with a result that relates to the survival function derived from the cumulative density function ($F(t)$) of the Weibull distribution:

$$S(t) = \mathrm{P}(T > t) = \int_t^{\infty} f(u)du = 1 - F(t) \quad [2]$$

which describes the proportion of surviving individuals (S(t)) at time t as:

$$S(t) = \exp\left\{-(d_0 \cdot t)^k\right\} \quad [3]$$

To fit the Weibull survival function to the data we used the log transformed form of the model:

$$\log_e(S(t)) = -(d_0 \cdot t)^k \quad [4]$$

The log-transformed form of the survival function was fit to the log-transformed proportion of surviving individuals using the Nelder-Mead method in the `mle2` function from the `bbmle v1.0.20` package in R (2). For each population, we fit the model using 90 combinations of initial parameter values and chose the optimal model based on Akaike information criterion. We used the same approach to fit the exponential survival function to the data and conducted a likelihood-ratio test. From the estimated Weibull parameters, we define the mean time to death of a cell, $\bar{T}_d$, as:

$$\bar{T}_d = d_0^{-1}\Gamma(1 + 1/k) \quad [5]$$

To estimate the standard error of $\bar{T}_d$ we used the delta method to estimate the variance of $\bar{T}_d$ (3). Because $\bar{T}_d$ was analyzed on a $\log_{10}$ scale, we performed the delta method on $\log_{10}$ transformed estimates of $\bar{T}_d$. We define the variance as:

$$\sigma^2_{\log_{10}\bar{T}_d} = \left\{ \begin{pmatrix} \frac{\partial \log_{10}\bar{T}_d}{\partial d_0} \\ \frac{\partial \log_{10}\bar{T}_d}{\partial k} \end{pmatrix}^T * \Sigma_{k,d_0} * \begin{pmatrix} \frac{\partial \log_{10}\bar{T}_d}{\partial d_0} \\ \frac{\partial \log_{10}\bar{T}_d}{\partial k} \end{pmatrix} \right\} \quad [6]$$

Where $\Sigma_{\hat{k},\hat{d_0}}$ is the variance covariance matrix of $d_0$ and $k$ that was estimated using `bbmle`. The partial derivatives are:

$$\frac{\partial \log_{10}\bar{T}_d}{\partial d_0} = \log_{10}(e)d_0 \quad [7a]$$

$$\begin{aligned} \frac{\partial \log_{10}\bar{T}_d}{\partial k} &= \log_{10}(e)(\Gamma(1 + 1/k))^{-1}\ \Gamma'(1 + 1/k) - k^{-2} \\ &= -k^{-2}\log_{10}(e)(\Gamma(1 + 1/k))^{-1}\Gamma(1 + 1/k)\,\psi_0(1 + 1/k) \\ &= -k^{-2}\ \log_{10}(e)\ \psi_0(1 + 1/k) \end{aligned} \quad [7b]$$

Where $\partial \log_{10}\bar{T}_d/\partial k$ was derived using the chain rule, $\psi_0$ is the polygamma function of order zero, and $e$ is Euler's number. The pooled variance ($\sigma^2_{p,\ \log_{10}\bar{T}_d}$) was calculated for each replicate within a given taxon using the following formula:

$$\sigma^2_{p,\log_{10}\bar{T}_d} = \frac{\sum_{i=1}^{k}(n_i - 1)\sigma^2_{i,\log_{10}\bar{T}_d}}{\sum_{i=1}^{k}(n_i - 1)} \quad [8]$$

Where $\sigma^2_{p,\log_{10}\bar{T}_d}$ represents the ith population. Standard errors were calculated as $SE_{p,\log_{10}\bar{T}_d} = \sigma_{p,\log_{10}\bar{T}_d}/\sqrt{n}$, where n is the number of biological replicates within a given taxon.

We use the Weibull parameters to estimate the time until population size estimates can no longer be obtained, making the population effectively extinct. We establish the critical proportion of surviving individuals for each replicate population as $S_{ext} = N(t_{ext})\ /N(0)$, Here $N(t_{ext})$ is the population size where on average we can sample only a single CFU given our experimental design and sampling regime, which is $N(t_{ext})$ = 1 CFU * 50 mL * 1 mL/ 0.1 mL = 500 cells. Using $S_{ext}$ as the quantile of interest, time until extinction ($T_{ext}$) is calculated as:

$$T_{ext} = d_0^{-1}(-\log_e(S_{ext})^{1/k}) \quad [9]$$

Using the Delta method again, we define the variance as

$$\sigma^2_{\log_{10} T_{ext}} = \left\{ \begin{pmatrix} \frac{\partial \log_{10} T_{ext}}{\partial d_0} \\ \frac{\partial \log_{10} T_{ext}}{\partial k} \end{pmatrix}^T * \Sigma_{k,d_0} * \begin{pmatrix} \frac{\partial \log_{10} T_{ext}}{\partial d_0} \\ \frac{\partial \log_{10} T_{ext}}{\partial k} \end{pmatrix} \right\} \quad [10]$$

And calculate the partial derivatives as:

$$\frac{\partial \log_{10} T_{ext}}{\partial d_0} = \log_{10}(e) d_0 \quad [11a]$$

$$\frac{\partial \log_{10} T_{ext}}{\partial k} = -\frac{\log_e(\log_e(1/S_{ext}))}{\log_e(10)k^2} = -\frac{\log_{10}(\log_e(1/S_{ext}))}{k^2} \quad [11b]$$

Pooled variances and standard errors were calculated as described above.

**Single-cell staining.** To identify dead cells, we used the nucleic-acid stain SYTOX Green (ex/em 504/523 nm), which is impermeable to intact microbial membranes due to its positive charge. To identify metabolically active cells we used CTC (5-cyano-2,3-ditolyl tetrazolium), a membrane-permeable dye that is colorless until reduced by the electron transport system (ETS) (4–6). Once it is reduced, CTC becomes a charged, red-fluorescing formazan precipitate salt (ex/em 488/630 nm) that functions as a redox molecule, the final electron acceptor in the ETS instead of oxygen (4) or alternate electron acceptors commonly used by anaerobic bacteria (5). To identify all cells we used DAPI (4',6-diamidino-2-phenylindole, ex/em 350/470), a membrane permeable dye that binds to DNA. Aliquots of $100\mu L$ were taken from ancestral cellular cultures at stationary phase and after 1,000 days. Thes aliquots were incubated in 1 mM of SYTOX Green for 15 min at 37 °C, 5 mM CTC for 30 min, and 18 $\mu$mol mL-1 DAPI for 15 min. The triple-stained samples were filtered onto a 25 mm, 0.2 $\mu$m black polycarbonate filter (Marine Manufacturing, Clinton Township, MI) using vacuum filtration (< 30 kPa) and mounted onto a 1.0 mm thick glass slide. Filters were fixed between the slide and a 0.15 mm thick glass coverslip with BacLight mounting oil to minimize background fluorescence. Differentially stained cells were counted using epifluorescence microscopy. We used a Zeiss Axioplan microscope with a mercury lamp equipped with a blue-light filter (BP365, FT395, LP397), a green-light filter (450-490, FT 510, LP520), and a custom filter that allowed us to view CTC while avoiding excitation and emission overlap with the other fluorochromes. This custom filter was a Chroma Technologies Acridine Orange/Di-8-ANEPPS filter (excitation 480/30x, BS 505DC, emission 620/60m). Cells from each slide were randomly surveyed by moving across the filter from left to right. A new image was collected in each new field of view. Images were collected with an IMI Tech IMC-3145FT digital camera. Ten images were randomly collected per slide with each filter set. We used the FIJI/Image J program to count the number of active, dormant, and dead cells from each image.

**Construction of ΔspoIIE mutant .** We chose to introduce a mutation in spoIIE, which controls cell division during sporulation but also activates sigma F, a transcription factor required for spore development. Construction of the ΔspoIIE mutant was performed on *B. subtilis* KBS0812 using a modified form of a previously described SPP1 transduction protocol (7). SPP1 lysate was made using the donor strain *B. subtilis* 168 ΔspoIIE BKE00640 obtained from the Bacillus Genetic Stock Center. Inoculation was performed in LB with 10mM $CaCl_2$ and $5\mu g$/mL chloramphenicol. Transduction was validated by PCR using the primers DAS11: 5' TAAGACACCGCCCTTTCACG 3' and DAS12: 5' AGCAGCCATCCGTTATCAGC 3'. The donor strain was used as a positive control and the recipient strain was used as a negative control. After two rounds of isolation streaking to remove phage, a transduced colony was grown in LB with 10mM $CaCl_2$, $1\mu g$/mL erythromycin, and $5\mu g$/mL chloramphenicol. Loss of sporulation was tested by growing the recipient strain in Difco Sporulation Media and testing for heat resistance. The antibiotic resistance cassette was removed from the recipient strain using a previously described approach (8).

**Metabolomics.** Output from GC/MS was analyzed by converting the proprietary `.D` files supplied by the GC/MS instrument into `mzXML` files using the `ProteoWizard msconvert` tool (9). `mzXML` files were then assembled into 3 batches: Amino Acid Standards, LTDE Media Samples, and All Files; before compressing each batch into a zip file and uploading the zipped GC/MS data to the `Workflow4Metabolomics` Galaxy server (10). GC/MS peaks were then deconvoluted using `metaMS.runGC v2.1` with default parameters (11). Identified peaks for unknown compounds along with their corresponding intensities and retention times were output in the resulting `peaktable.tsv` file. Unknowns were annotated by matching the associated mass spectra provided in the `peakspectra.msp` output with characterized metabolites in the GOLM Metabolome database using the ms analysis tool with no GC column-type or retention index selected (12). Finally, to confirm the presence and identity of annotated peaks, we additionally used the Quantitative Analysis tool from the instrument supplied, proprietary Mass Hunter (Aglient) software. Metabolite identities of associated mass spectra were annotated by querying the NIST11 database.

Targeted metabolomics was performed on *Bacillus* sp. KBS0812 cell-free supernatant using a previously described approach (13). Targeted metabolomics was performed on each *Bacillus* sp. KBS0812 replicate population using three technical replicates. Blank measurements were subtracted from the measurements obtained for each sample.

**Phylogenetic reconstruction.** PCR on the 16S rRNA gene of each strain was performed using 8F and 1492R primers and PCR reaction conditions previously described (14). PCR products were purified using the QIAGEN QIAquick PCR Purification Kit and Sanger sequenced at the Indiana Molecular Biology Institute (IMBI) at Indiana University Bloomington (IUB). Sequences were aligned using SILVA INcremental Aligner v1.2.11 (`SINA`) (15). Phylogenetic reconstruction was performed using of Randomized Axelerated Maximum Likelihood v8.2.11 ( `RAxML`) (16). The phylogeny was inferred using the General Time Reversible (GTR) model of nucleotide substitution with gamma distributed rate variation. Bootstrap convergence criteria were set to `autoMRE`. Rapid bootstrap analysis and the search for the best-scoring ML tree was performed in a single program call. The 16S rRNA sequence of *Prochlorococcus marinus* subsp. marinus str. CCMP1375 (NCBI accession number NC_005042) was used as an outgroup.

**Modeling trait evolution.** We modeled the evolution of $d_0$ on a rooted ultrametric form of our 16S rRNA RAxML phylogeny. Phylogenetic comparisons were performed using the Phylogenetic Monte Carlo (`pmc`) package `v1.0.3` (17). Pairwise model comparisons were performed for Brownian motion vs. Pagel's lambda using 1,000 iterations. We removed *Bacillus* sp. KBS0812 from our data for this analysis, as it represented an extreme observation with a phylum-specific trait that the remaining taxa do not have. All statistical analyses were performed in `R v3.5.0` .

**Genome sequencing and assembly.** We performed whole genome sequencing using two sequencing technologies to obtain contiguous reference genomes for each taxon. Purified DNA was prepared for sequencing using the Illumina TruSeq DNA sample prep kit with an insert size of 250 bp and sequenced on an Illumina HiSeq 2500 using 100 bp pair-end reads (Illumina, San Diego, CA) at the Center for Genomics and Bioinformatics (CGB) at Indiana University Bloomington (IUB) for the following strain designations: KBS0701, KBS0702, KBS0703, KBS0705, KBS0706, KBS0710, KBS0711, KBS0713, KBS0714, KBS0715, KBS0721, KBS0722, KBS0724, KBS0725, KBS0727, KBS0801, KBS0802. DNA libraries were constructed using the Nextera DNA Sample Preparation kit with an insert size 300 on the Illumina HiSeq 2500 using 300 bp pair-end reads (Illumina, San Diego, CA) at the Hubbard Center for Genome Studies, University of New Hampshire for the following strain designations: ATCC13985, ATCC43928, KBS0702, KBS0707, KBS0712, KBS0801, KBS0812. Raw FASTQ reads were processed by removing Illumina TruSeq adaptors, trimming the end of each read, and quality-filtering for an average Phred score of 30 using `cutadapt v1.7.1` (18).

DNA extraction for Nanopore sequencing was performed using a previously described method (19). Library preparation was performed using the Nanopore Ligation Sequencing Kit (SQK-LSK109) and the 1D native barcoding genomic DNA procedure using barcode kit EXP-NBD104 and library kit SQK-LSK109. MinION sequencing was performed using the manufacturer's guidelines on `R9.4.1` flow cells (FLO-MIN106) on a MinION `18.12.9` (Oxford Nanopore Technologies, Oxford, United Kingdom). Base-calling and de-barcoding was performed using `Guppy v2.3.5` with configuration files `dna_r9.4.1_450bps.cfg` and `configuration.cfg`. We kept reads longer than 1,000 bp with an average Phred quality score of at least 10 and cut the first 100 bp using `NanoFilt v2.3.0` (20). Hybrid assemblies were generated using `Unicycler v0.4.7` (21).

**Comparative genomics.** Because our set of taxa is phylogenetically diverse they have few orthologues, making it difficult to determine whether convergent evolution occurred at the gene level.

The metabolic pathway composition was inferred using the Metabolic And Physiological potentiaL Evaluator (`MAPLE v2.3.1`; (22)). MAPLE was run using bi-directional best hit with NCBI BLAST on KEGG genes and modules version `20190318` using all prokaryotes in KEGG. MAPLE output files for the module pathways, signatures, and complexes were all filtered for query coverage values greater than 80% and merged into a single file for each taxon. Filtered MAPLE results for were merged into a single presence-absence matrix.

**Mutation calling.** Genome-wide pairwise nucleotide diversity was estimated from mutations called as SNPs by `Breseq v0.32.0` (23). We found no evidence of fixed mutations in any population. The few reasonable candidates that `Breseq` classified as fixed had extremely low coverage ($\leq 10$), suggesting that they were unlikely to represent "true" fixations and were removed from downstream analyses.

Because the ancestral population was grown from a single CFU for only $\sim 13$ generations, the high number of detectable mutations and their frequencies cannot be explained by the presence of ancestral genetic variation. Rather, these frequencies are the result of *de novo* mutations and birth events that occurred over 1,000 days of energy-limitation. However, to be conservative we chose to only examine mutations that were observed within a single replicate. Estimates of genetic diversity was calculated for all SNPs across the genome, as we assumed that recombination is rare and most sites are effectively linked. Nucleotide diversity was calculated using the bi-allelic equivalent of the pair-wise formula:

$$\hat{\theta}_{\Pi} = \frac{n_c}{n_c - 1} \frac{1}{L} \sum_{i} 2 * p_i \, (1 - p_i) \qquad [12]$$

Where $p_i$ represents the frequency of the $i$th mutant and the term $n_c$ represents the number of chromosomes in the pooled library. Since we sequenced the population from a bulk DNA extract, we chose to use the final population size as $n_c$. We calculated the sample size-corrected number of segregating sites ($\hat{\theta}_W$) and Tajima's D ($T_D$) as previously described (Gillespie, 2004). The ratio of non-synonymous to synonymous mutations ($pN/pS$), genome-wide likelihood ratios, and gene-specific multiplicity scores were calculated as previously described (24). We tested whether $pN/pS$ was less than one in each taxon using a left-tailed one-sided $t$-test. Using mean allele frequencies ($\bar{f}$) and assuming that mutant lineages grew as a binary tree, we estimated the number of generations ($t_m$) as $\lfloor log_2(\bar{f} \cdot N(t_{final})) \rfloor$ and the number of cell divisions as $\sum_{i=1}^{t_m} 2^i$ for all replicate populations where we could call mutations. Linear mixed models were fit using `statsmodels 0.11.1` (25).

Index of replication (iRep) values were estimated in `Python` using the `iRep` package `v1.1.14` (26). The `iRep` code was run on Sequence Alignment Map (SAM) files that were mapped using `BWA-MEM v0.7.12` and converted to SAM format using `SAMtools v1.9` (27).

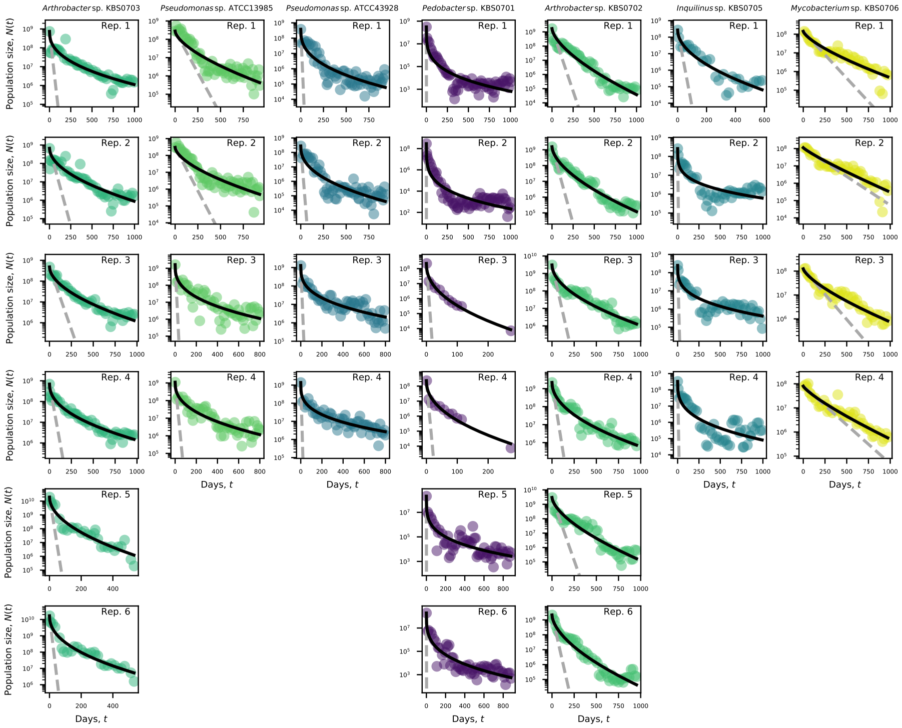


**Fig. S1.** The survival curve of each replicate population across taxa.

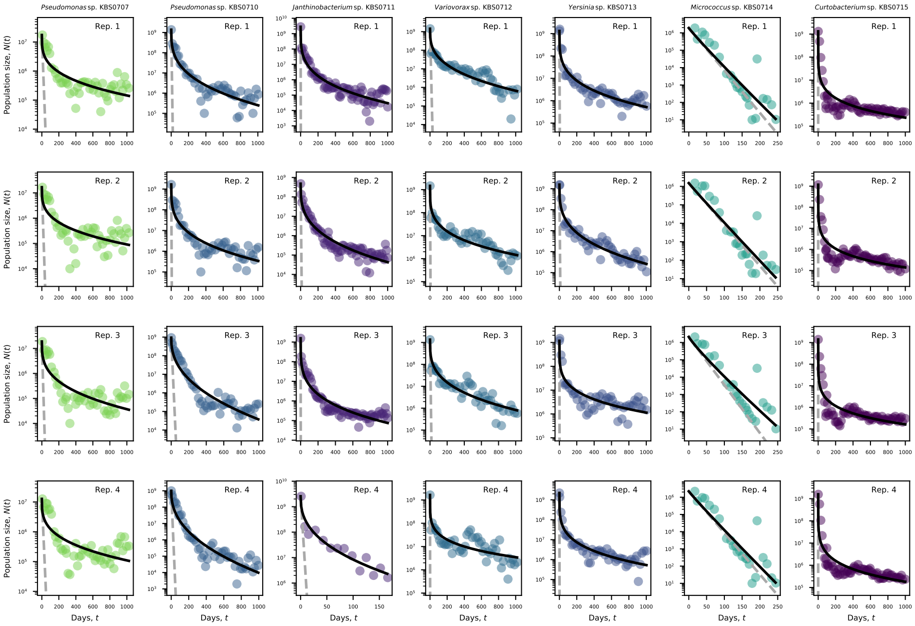


**Fig. S2.** The survival curve of each replicate population across taxa.

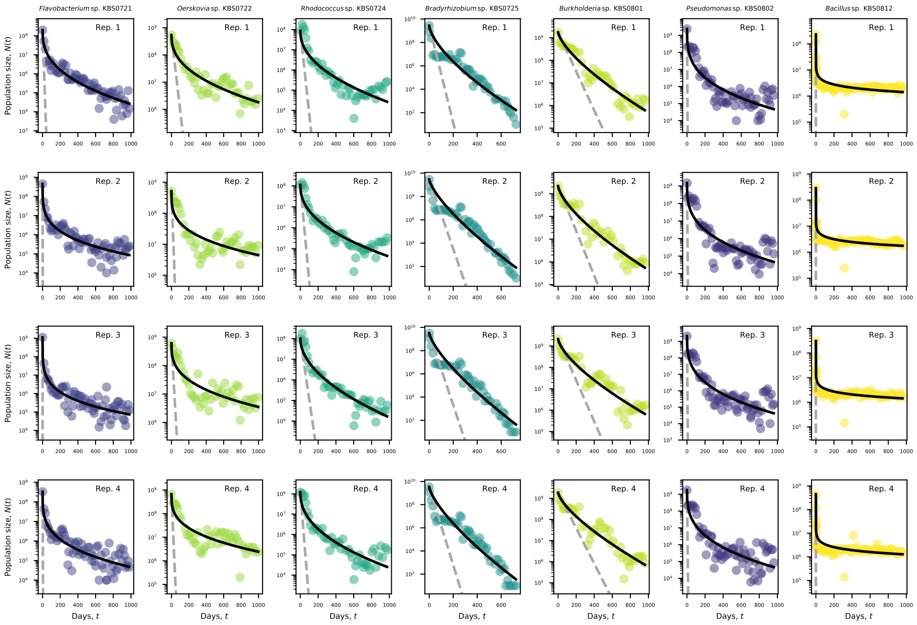


**Fig. S3.** The survival curve of each replicate population across taxa.

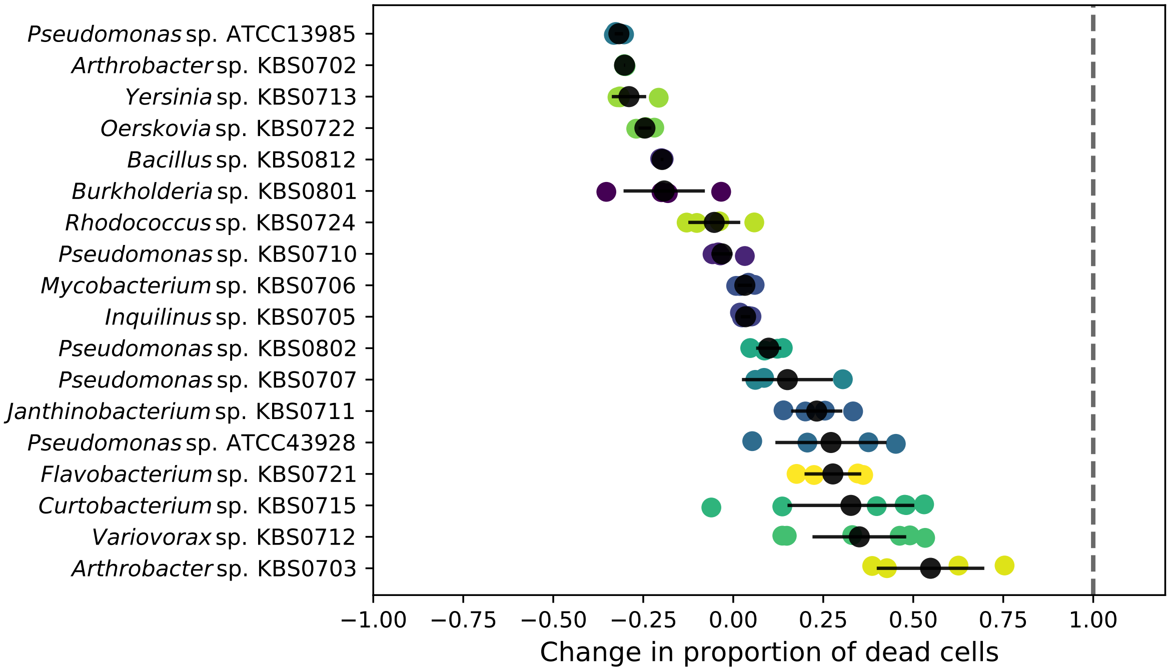


**Fig. S4.** The change in the proportion of dead cells across taxa. The black dot represents the mean change and the black bars represent twice the standard error.

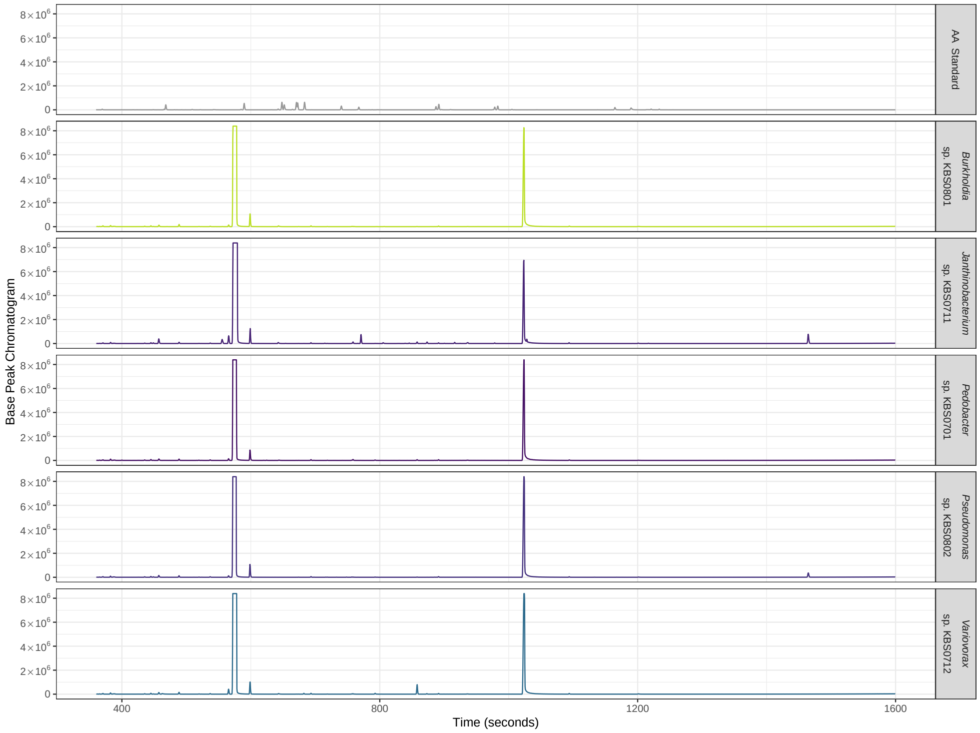


**Fig. S5.** Chromatogram summarizing the mass spectrometry profiles of the cell-free supernatant of five taxa at day 1,000. The figure in the top row is the amino acid standard. The three highest peaks in plots of taxa are internal standards. The concentration of amino acids is below the detection limit in all samples.

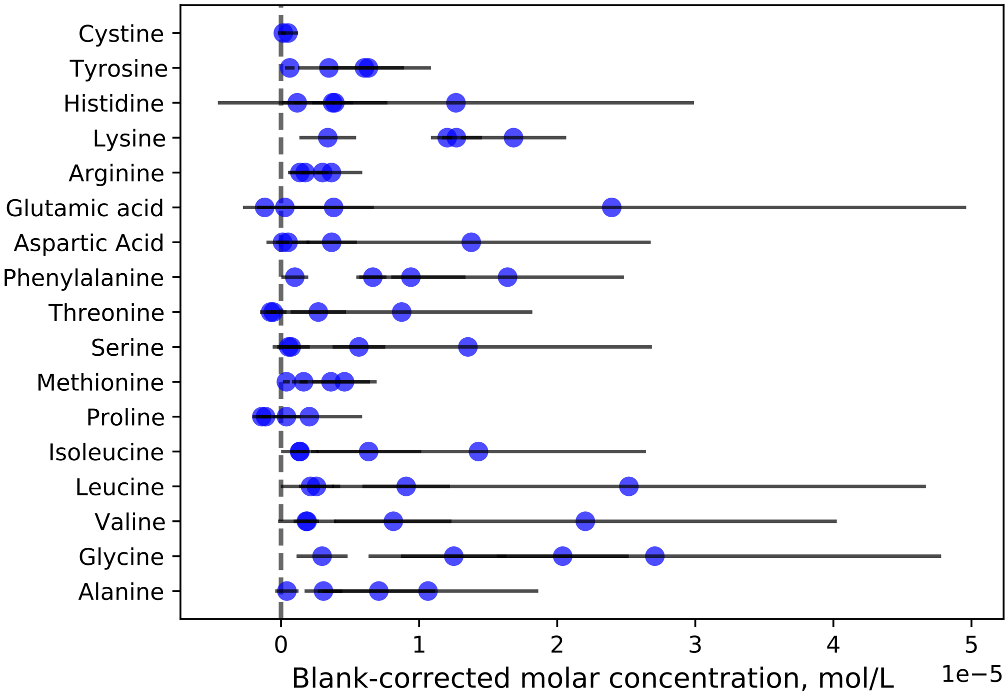


**Fig. S6.** Molar concentrations of amino acids in cell free supernatant of $B.\ subtilis$ KBS0812. Each dot and bar represents the mean and twice the standard error for a given replicate population, respectively. These concentrations are on the order of the blank, despite the fact cell death estimates were on the order of $10^6 - 10^7\ \mathrm{mL}^{-1}$. Targeted metabolomics was performed as previously described (13).

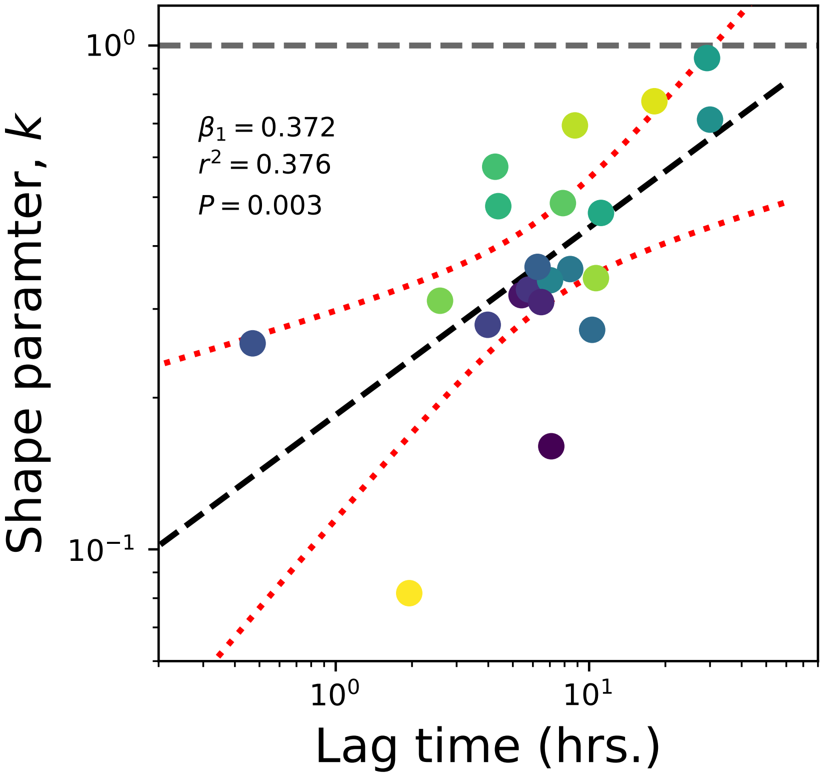


**Fig. S7.** There is a strong linear relationship between Lag time and the shape parameter of the Weibull distribution. Each taxon is represented by a dot. The dashed black line is the slope of a simple linear regression. The red dotted lines represent the 95% confidence hull. The dashed grey horizontal line indicates a shape parameter value of one, where the Weibull reduces to an exponential.

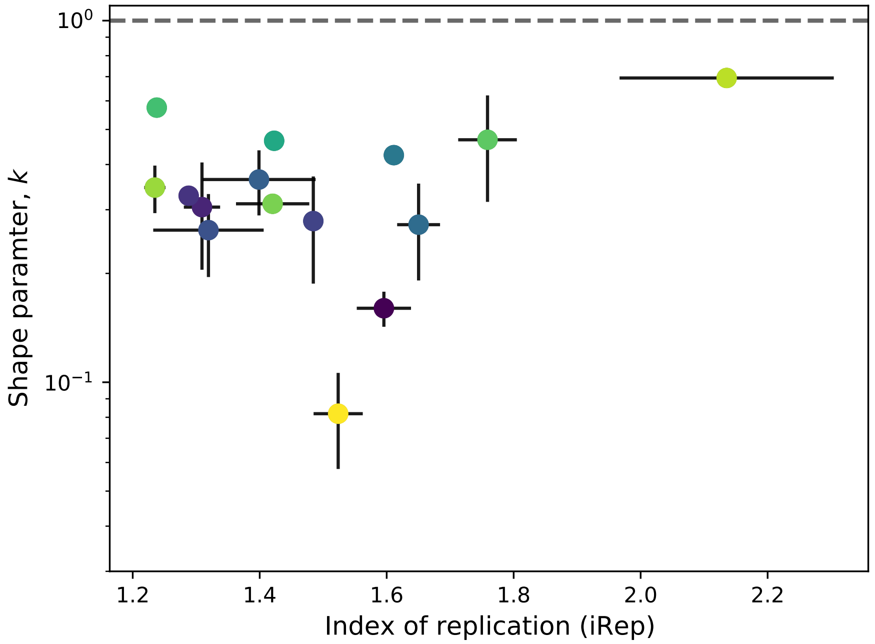


**Fig. S8.** A scatterplot of iRep and the shape parameter of the Weibull distribution. Values are plotted for all samples with sufficient sequence coverage to estimate iRep (described in methods). Black dots represent mean values and black lines represent twice the standard error of mean. There is no visible relationship and no the slope of a mixed effect linear model with random slopes was not significant. The dashed grey horizontal line indicates a shape parameter value of one, where the Weibull reduces to an exponential.

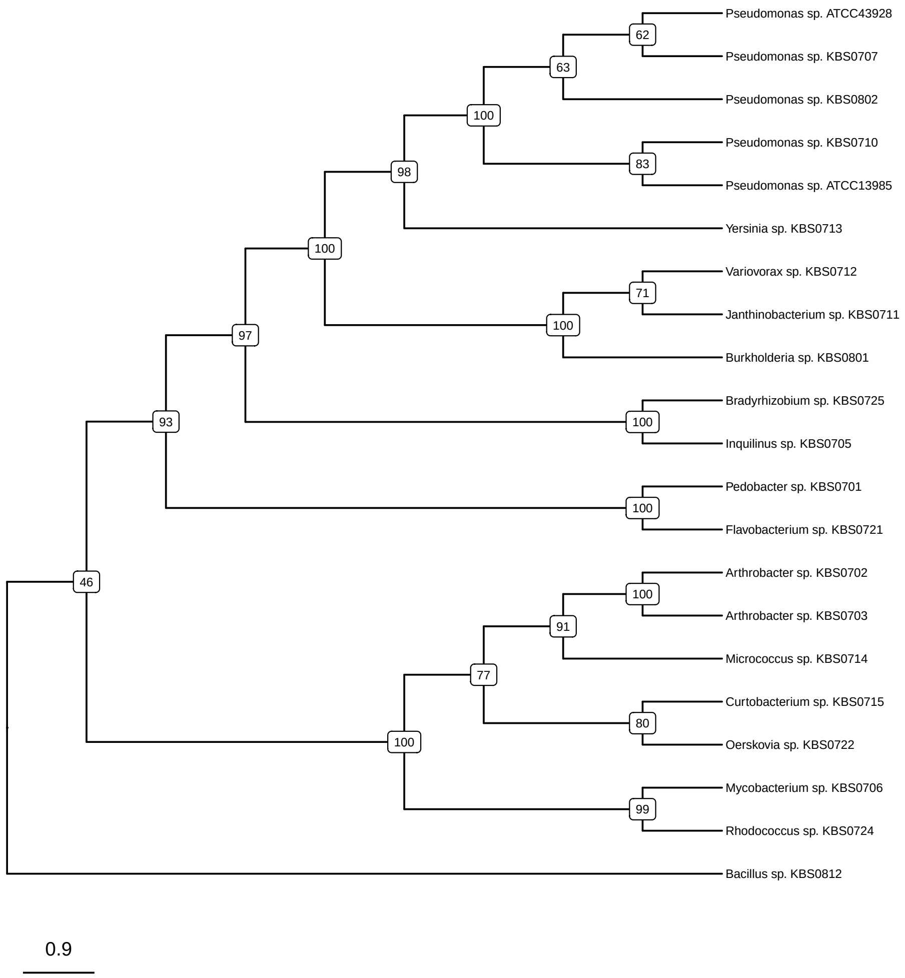


**Fig. S9.** The ultrametric RAxML 16S rRNA phylogenetic tree of the taxa used in this study. Numbers represent bootstrap support values. Outgroup not shown.

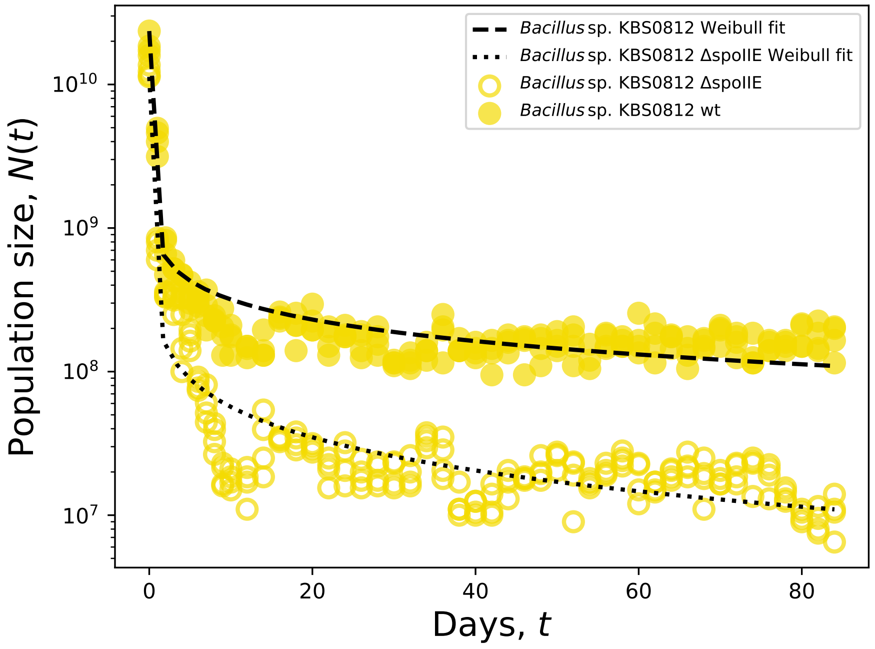


**Fig. S10.** The survival curves of replicate populations of *Bacillus* sp. KBS0812 and its ΔspoIIe mutant show that the ability to form protective endospores does not affect the survival curve over a timescale of 20 days. The solid black and dashed grey lines indicate the fit from the survival function of the Weibull and exponential distributions, respectively.

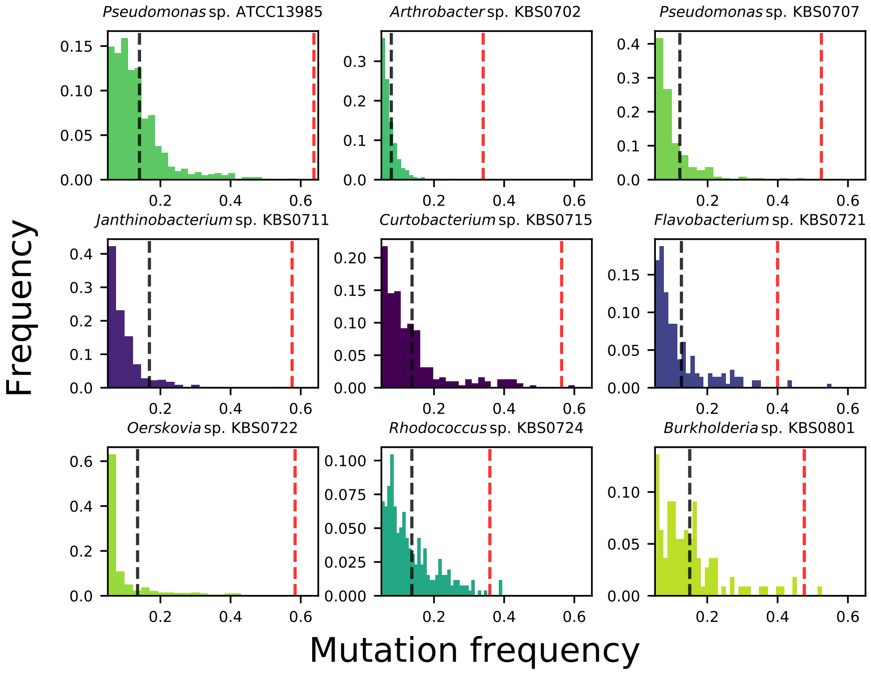


**Fig. S11.** The site frequency spectra of all taxa that meet our filtering criteria. The dashed black line represents the mean of the mean mutation frequency and the dashed red line represents the mean of the maximum observed mutation frequency across replicate populations, respectively.

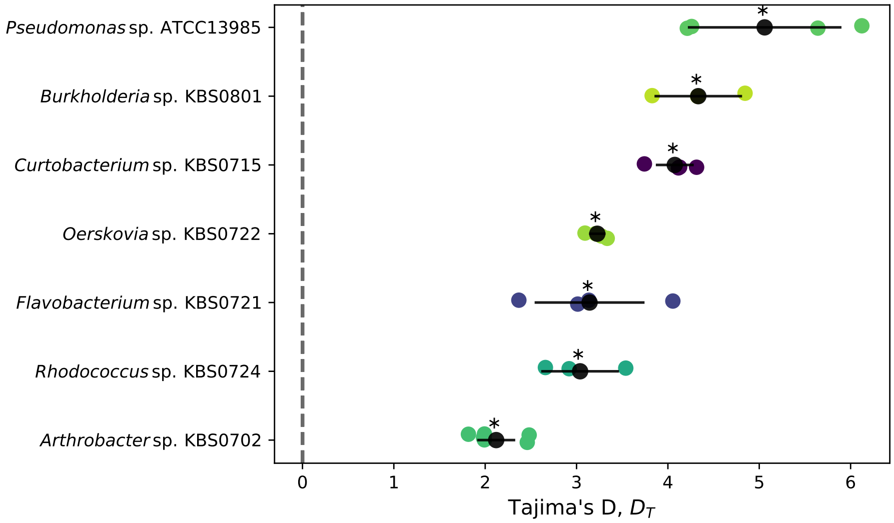


**Fig. S12.** Tajima's D ($D_T$) values for all taxa with a sufficient number of mutations in at least three replicate populations. The point where the mean number of pairwise differences is equal to the number of segregating sites in the population is represented by a dashed grey vertical line. The black dot represents the mean $D_T$ within a given taxon and the black bars represent twice the standard error. The asterisk indicates that $D_T$ is significantly greater than zero using a right-tailed one-sided $t$-test given a false discovery rate of 0.05.

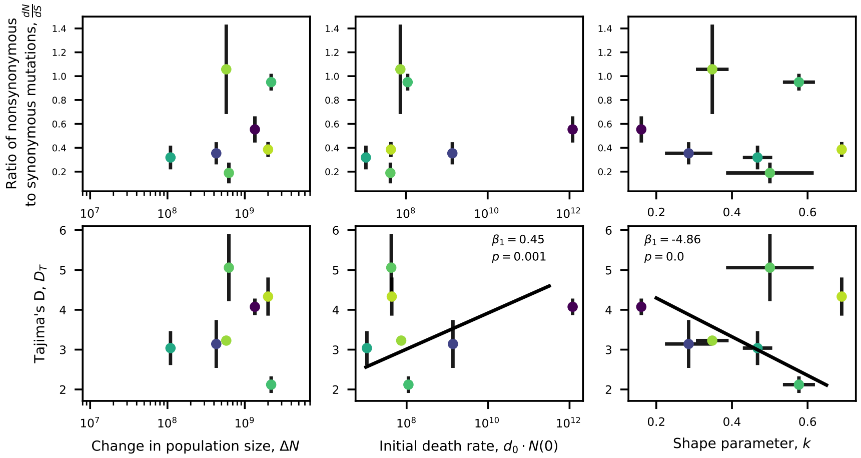


**Fig. S13.** There is no clear relationship between $dN/dS$ and $D_T$ and measures of demography across taxa. Slopes were not significant for mixed linear models with random taxon-specific intercepts for seven out of eight relationships. A borderline significant relationship was found between $k$ and $d_0 \cdot N(0)$ and $D_T$.

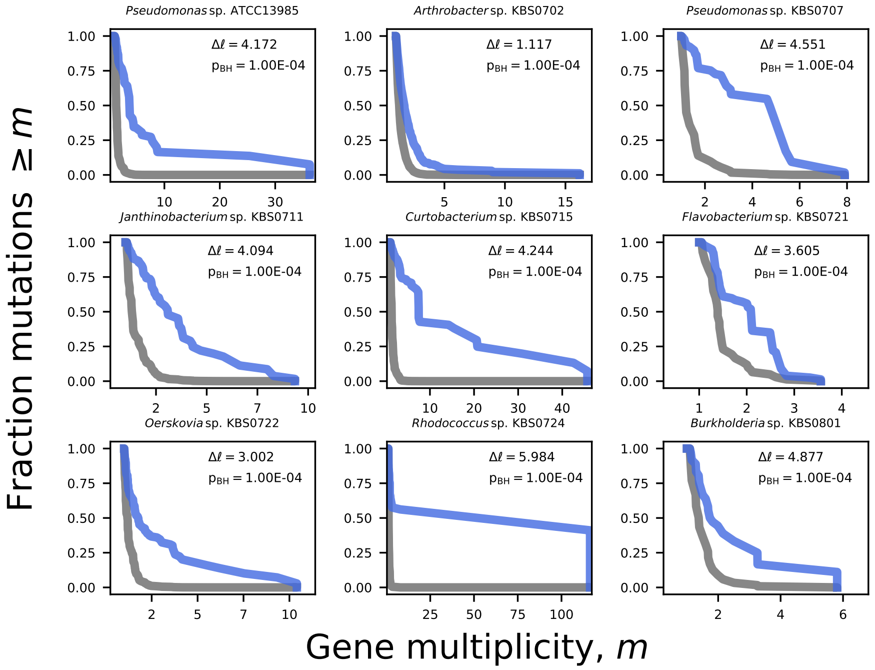


**Fig. S14.** Survival curves of the number of non-synonymous mutations observed within a gene, normalized by relative gene length (i.e., multiplicity, $m$). We find that genes are significantly more enriched for mutations than expected by chance in all taxa that meet our criteria (SI Appendix; Table 1). The grey line represents the null expectation (eq. 72 from the SI of (24)). The genome-wide net increase in the log-likelihood of an excess of mutations relative to the null model ($\Delta\ell$) and its Benjamini-Hochberg corrected $p$-value is included in each sub-plot (eq. 74 from the SI of (24)).

### SI Dataset S1 (weibull_results_clean_species.csv)

Summary statistics for the mean survival curve results of all taxa. Each column represents the mean value of that variable across replicates of a given taxon.

### SI Dataset S2 (total_parallelism.txt)

Genome-wide parallelism scores for all taxa that acquired at least 50 non-synonymous mutations across all replicates.

### SI Dataset S3 (gene_annotation.txt)

The RefSeq annotations of all significantly significant genes for all taxa with their respective annotated function.

### SI Dataset S4 (genomes_info.txt)

Annotation information for all reference genomes assembled for this study.
